# Computational modeling of virucidal inhibition mechanism for broad-spectrum antiviral nanoparticles and HPV16 capsid segments

**DOI:** 10.1101/2021.08.21.457236

**Authors:** Parth Chaturvedi, Payam Kelich, Tara A. Nitka, Lela Vuković

## Abstract

Solid core nanoparticles coated with sulfonated ligands that mimic heparan sulfate proteoglycans (HSPG) can exhibit virucidal activity against many viruses that utilize HSPG interactions with host cells for the initial stages of the infection. How the interactions of these nanoparticles with large capsid segments of HSPG-interacting viruses lead to their virucidal activity has been unclear. Here, we describe the interactions between sulfonated nanoparticles and segments of the human papilloma virus type 16 (HPV16) capsids using atomistic molecular dynamics simulations. The simulations demonstrate that nanoparticles primarily bind at interfaces of two HPV16 capsid proteins. Insertions of nanoparticles at these interfaces leads to increased separation in distances and angles between capsid proteins. As the time progresses, the nanoparticle binding can lead to breaking of contacts between two neighboring proteins. The revealed mechanism of nanoparticles targeting the interfaces between pairs of capsid proteins can be utilized for designing new generations of virucidal materials and contribute to the development of new broad-spectrum non-toxic virucidal materials.

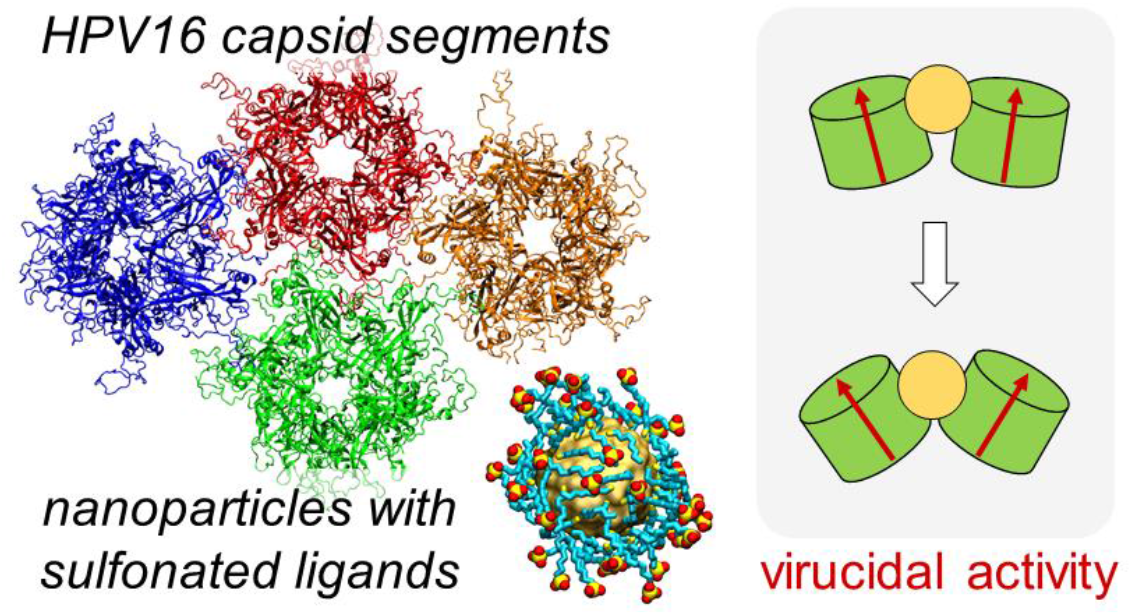

## 1. Introduction

Among many infectious disease threats that humans face from microorganisms, viral infections are arguably the biggest pandemic threat at present. Even in the absence of pandemics, viral infections kill millions of people every year, as viruses have high rates of replication, mutation and transmissibility^1^. Available antiviral drugs are often highly targeted against specific viruses or, in some cases, members of a viral family^2^. Current antiviral drugs include small molecules, such as nucleoside analogues and peptidomimetics, monoclonal antibodies, proteins that are able to stimulate the immune response, such as interferon, and oligonucleotides^2^. Many of these drugs act intracellularly with selectivity for viral enzymes. However, since viruses mostly depend for their replication on infected host cells, the specificity of antiviral drugs for viral proteins is not ideal, often causing general intrinsic toxicity upon administration^3,4^. Furthermore, due to high mutation rates, most viruses develop drug resistance^5– 7^.

When antiviral drug development is based on single virus-specific protein as a target, the discovered therapeutics lack the broad-spectrum effects and thus the capability of targeting many viruses that are phylogenetically unrelated and structurally different. Yet, there is a demand for development of broad-spectrum antiviral therapeutics that are non-toxic to hosts. One development approach is to identify substances that can favorably interact with many viruses outside of host cells and determine if these interactions can prevent the first stages of infection and the viral replication cycle. Previous efforts, based on the principle of targeting virus-cell interactions that are common to many viruses, identified many non-toxic substances that interact with a broad spectrum of viruses, such as heparin and polyanions^8–12^. However, most of these substances exhibit only virustatic properties where their activity depends on a reversible binding event. Binding reversibility makes these substances therapeutically ineffective: upon dilution, these materials detach from intact viral particles allowing the viruses to infect again.

Virucidal materials show large promise for use against viral infections. Virucidal molecules cause irreversible viral deactivation, which is not adversely affected by dilution^13^. Virucidal materials can include detergents, strong acids, polymers, and nanoparticles^14–18^. However, most of these materials have intrinsic cellular toxicity since they attempt to chemically damage the virus and they often cannot selectively damage the virus without affecting the host in which the virus replicates.

Recent studies demonstrated that non-toxic broad-spectrum virucidal materials can be developed based on the principle of mimicking the virus-host interactions common to many viruses^19,20^. In particular, virucidal material design can be inspired by viral attachment ligands / host cell receptors called heparan sulfate proteoglycans (HSPG), which are responsible for the initial steps of the virus replication cycle^21^. Many viruses, including human papillomavirus type 16 (HPV-16), human immunodeficiency virus 1 (HIV-1), herpex simplex virus 1 and 2 (HSV-1 and HSV-2), attach to HSPGs, which are expressed on the surfaces of almost all eukaryotic cell types^22^. Nanoparticles (NPs) coated with long and flexible sulfonated ligands mimicking HSPG were shown to effectively associate with such viruses and eventually lead to irreversible viral deformation. These non-toxic nanoparticles (NPs) show *in vitro* irreversible virucidal activity against HSV-2, HPV-16, RSV, Dengue and lentivirus, and are also active *ex vivo* in human cervicovaginal histocultures infected by HSV-2 and *in vivo* in mice infected with RSV^19^.

Previous transmission electron microscopy analyses suggested the mechanism of the virucidal activity of NPs coated with sulfonated ligands against the HPV16 virus: NPs eventually break the capsid or the virus particle^19^. In the present study, we use large scale atomistic molecular dynamics (MD) simulations to explore this suggested mechanism on systems containing 2.4 nm gold core nanoparticles coated with HSPG-mimicking ligands and extended capsid segments of two, three and four HPV16 major late (L1) pentamer proteins. Our studies expand on previous modeling of sulfonated nanoparticles interacting with *single* HPV16 capsid proteins^19^. The present simulations identify the preferred sites of interaction between sulfonated NPs and capsid segments, as well as the perturbations induced in the capsid segments by the presence of NPs.

## 2. Methods

### Constructing HPV16 Capsid Segments

The cryo-electron microscopy (cryo-EM) structure of the Human Papillomavirus Type 16 (HPV16) L1 capsid proteins, assembled into a capsid structure, shown in **Figure 1a**, was obtained from the RCSB protein databank (PDB: 3J6R)^23^. Segments of either two, three or four L1 proteins with the least empty space between them were selected and then extracted from the capsid. As L1 protein structures in the extracted segments were defined via the coordinates of the backbone atoms only, each L1 protein structure in three extracted segments was replaced by a crystal structure of HPV16 L1 protein (PDB: 5W1O)^24^. In all three segments, each L1 pentamer structure was docked into the cryo-EM structure-based protein density using *colores* tool in Situs software using a resolution of 5 Å^25^. Structures of resulting three HPV16 capsid segments were then further flexibly fitted into cryo-EM-extracted densities of L1 proteins using the molecular dynamics flexible fitting (MDFF) method in NAMD 2.13^26–28^. MDFF simulations, using the CHARMM36 force field^29^, were carried out for 50,000 steps with timestep of 1 fs in the constant volume and temperature conditions at 310 K in vacuum, with the scaling factor for potential, g_scale_ (ξ), set to the value of 0.3 and with Langevin constant γ_Lang_ set to 5.0 ps^−1^. Secondary structure, cispeptide bonds, and chirality restraints were applied on L1 pentamers to prevent the unnatural distortions during the fitting procedure.

**Figure 1.**
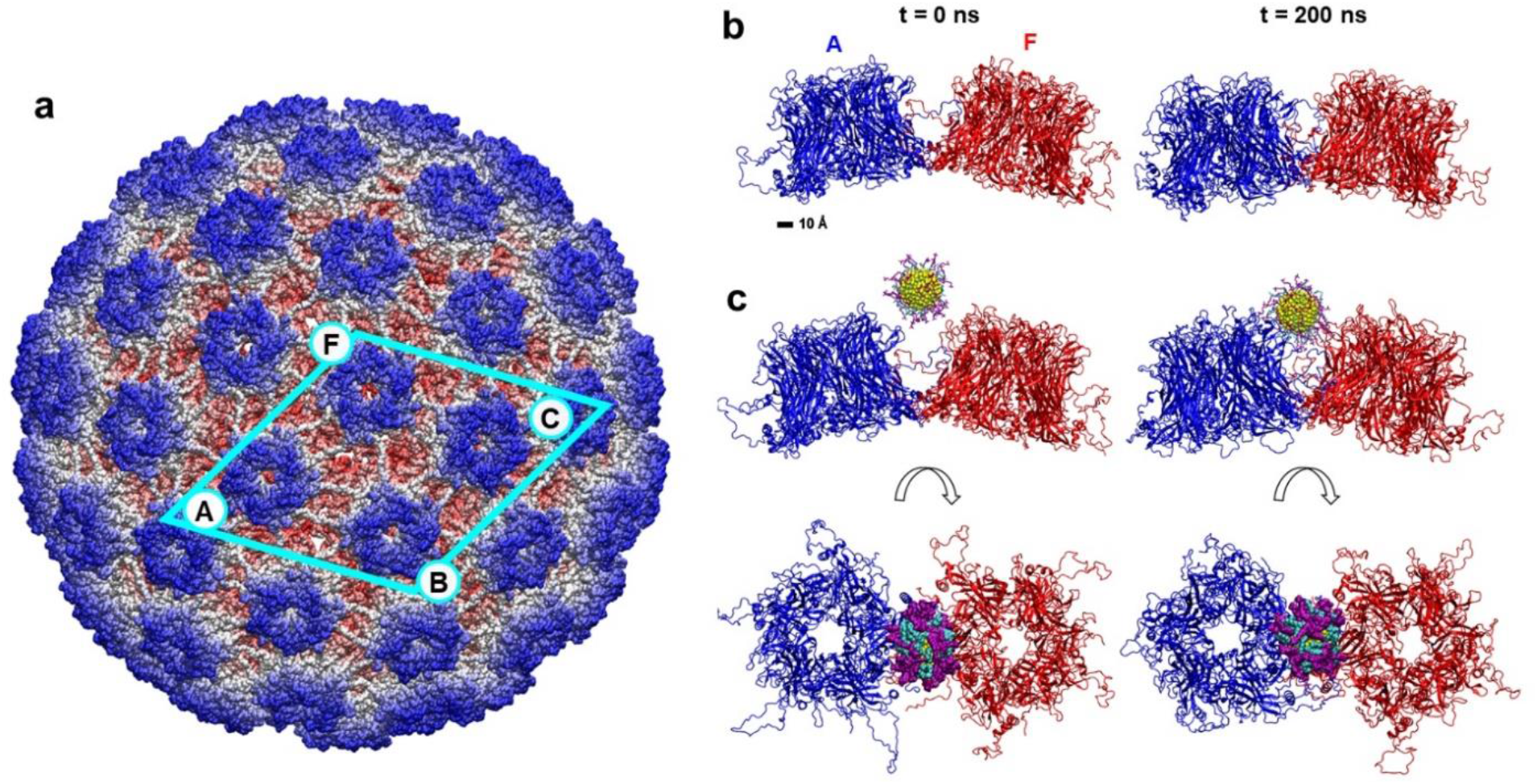
HPV16 virus capsid and its segments formed by L1 pentamer proteins. a) Cryo-EM structure of the HPV16 capsid, assembled from L1 pentamer proteins. Labeled pentamers contribute to capsid segments examined in MD simulations. The largest segment examined, formed by four L1 pentamers, is marked by the cyan frame. The atoms are colored according to their radial distance from the capsid center. Pentamer labels (A, F, B, C) correspond to labels used in figures and analyses described below. b) Conformations of the capsid segment containing two L1 pentamers at the initial time and after 200 ns of MD simulations. The pentamers are colored in blue (A) and red (F), and the aqueous solution is not shown for clarity. c) Conformations of the same segment in the presence of the MUS:OT gold nanoparticle at the initial time (left) and after 200 ns of MD simulations (right), presented in the side and top views. Gold atoms, MUS ligands and OT ligands are shown in yellow, purple, and teal, respectively.

Individual chains of L1 protein crystal structures fitted into capsid segments were missing single loop regions and several residues at C- and N-terminals. Therefore, after MDFF fitting, the missing residues were modeled as flexible coils and covalently attached to the full protein, to provide complete L1 protein structures within capsid segments. Disulfide bonds were built between C161 and C324 residues for all the chains in L1 proteins. Final completed capsid segments were solvated with TIP3P water and neutralized with 0.15 M NaCl with *solvate* and *ionize* VMD plugins^30^, respectively.

### Ligand Docking on HPV Pentamer

Small sulfonated ligands (CH_3_CH_2_SO_3_^-^) were docked onto a single L1 protein and a pair of L1 proteins using the Autodock Vina software^31,32^. Docking was performed using the finalized structures of one and two L1 proteins described above. Sulfonated ligand structures were built using AMPAC graphical user interface. In Autodock Vina, the grid box was centered at various positions on a grid, scanning the protein surfaces with five fixed z-coordinates and changing the x and y coordinates from -40 Å to 40 Å for both, with an interval of 1 nm. Docking procedure was automated using our in-house Linux shell script. Each grid box dimension was 1 × 1 × 1 nm^3^ with a default spacing and exhaustiveness of 0.0375 nm and 8, respectively. The above procedure determined the locations of the ligands with the lowest docking scores on the L1 protein surfaces (**Figure S1**). These locations guided the placement of the nanoparticles with sulfonated ligands above the capsid segments.

### Nanoparticle Model

Atomistic model of a spherical nanoparticle was prepared by cutting a face centered cubic lattice of gold (Au) atoms (Au-Au bond length 2.88 Å) into a sphere with diameter of ∼2.4 nm. The gold core was ligated with a random spherical array of 40 1-octanethiol (OT) and 40 11-mercapto-1-undecanesulfonate (MUS) ligands. All ligands were built with AMPAC, and then arranged into a random spherical array with 0.3949 per Au atom surface density using our own code. The built NP was then solvated in TIP3P water and ionized with sodium (Na^+^) and chloride (Cl^-^) ions at a 0.15 M NaCl concentration using the VMD *solvate* and *ionize* plugins^30^. After solvation and ionization, short MD simulations were performed to obtain a structure of the equilibrated NP as an input for the NP-capsid segment simulations. NPs, described with the generalized CHARMM force field^33^, were minimized for 2,000 steps using NAMD2.13 software^26^ and equilibrated for 20 ns in the NPT ensemble with Langevin dynamics (Langevin constant γ_Lang_ = 1.0 ps^−1^), where temperature and pressure remained constant at 310 K and 1 bar, respectively. The particle-mesh Ewald (PME) method was used to calculate the Coulomb interaction energies, with periodic boundary conditions applied in all directions. Evaluation of van der Waals and Coulombic interactions was performed every 1- and 2-timesteps, respectively. A timestep of 2 fs was used for equilibration of NPs.

### Molecular Dynamics Simulations

Atomistic simulations were conducted to investigate interactions of HPV16 capsid segments alone and in the presence of a 2.4 nm-core MUS:OT NP. All the systems were described with CHARMM36 force field parameters^29,33^. MD simulations were performed with the NAMD2.13 package^26^. All simulations were conducted with Langevin dynamics (Langevin constant γ_Lang_ = 1.0 ps^−1^) in the NPT ensemble, where temperature and pressure remained constant at 310 K and 1 bar, respectively. The particle-mesh Ewald (PME) method was used to calculate the Coulomb interaction energies, with periodic boundary conditions applied in all directions. The evaluation of van der Waals and Coulombic interactions was performed every 1- and 2-timesteps, respectively.

In total, six systems were prepared for final MD simulation runs. They consisted of two, three and four HPV16 L1 pentamers with and without the nanoparticle. Guided by docking results, systems with nanoparticles were prepared by placing a single nanoparticle 10 Å above the interface of three pentamers or two pentamers in case of a two pentamer segment. For all systems, 10,000 steps of minimization were performed, followed by 2 ns equilibration of solvent molecules around the pentamers, which were restrained by using harmonic forces with a spring constant of 1 kcal/(mol Å^2^). Next, the systems were equilibrated in production MD runs, with center-of-mass (COM) restraints applied to α-carbons of a buried residue (chosen to be residue 331) for all five chains of L1 pentamers with a force constant of 2.0 kcal/mol, to keep protein molecules in the original unit cell. To prevent excessive rotations of pentamers in space, which, if allowed, would require much larger sizes of the water box, an additional 10 Å-wide square potential along one of the system axes was applied on L1 pentamers, using the wall potential constant of 100 kcal/mol.

### Data Analysis

#### Distance between two pentamers

To analyze distances between two L1 pentamers, we calculated distances between centers of mass (COM) of backbone atoms of individual pentamers over the duration of simulations using our scripts in VMD^30^.

#### Angle between two pentamers

To analyze tilting of two pentamers with respect to each other, we calculated angles between vectors that align with axes of the cylindrically shaped void space in pentamer centers. Directions of these vectors were calculated by approximating shapes of pentamers as cylinders and evaluating the direction of the plane passing through COM of atoms of every alternating monomer (three in total) that forms each pentamer protein.

#### Residue Contacts

To analyze interaction between neighboring pentamers and between NP and pentamers, we evaluated contacts between different protein residues and between protein residues and NP over the duration of simulations. A contact was defined to exist at a given time frame if any atom of the concerned protein residue was located within 5 Å of the other pentamer or the NP.

## 3. Results and Discussion

The present study examines the interactions between gold nanoparticles coated with sulfonated ligands and segments of HPV16 capsids of varying sizes. Previous modeling of these virucidal NPs interacting with a single L1 capsid protein of the HPV16 virus suggested that the initial steps of the virucidal mechanism involve NPs binding multivalently to L1 proteins via charged, polar and hydrophobic interactions^19^. However, the atomistic details behind the NP interactions with larger HPV16 capsid segments and the eventual distortion and breaking of the capsid, proposed to be at the core of the virucidal activity, remained unexamined and unclear. Here, we probe the effects of the NP binding on the stability of the HPV16 capsid segments. We first modeled several segments of HPV16 capsid and relaxed them in MD simulations. Then, these segments were also simulated in the presence of virucidal MUS:OT NPs. Two sets of simulations were then analyzed to determine the mechanism of virucidal activity by these nanoparticles.

### MD Simulations of HPV16 Capsid Segments in the Presence and Absence of Nanoparticles

A total of six systems, containing segments of two, three and four L1 proteins by themselves and in the presence of single MUS:OT nanoparticles, were modeled in aqueous solution and examined in atomistic MD simulations. The snapshots of these systems after 200 ns (for two- and three-pentamer systems) and 115 ns (for four-pentamer systems) of equilibration are shown in **Figures 1** and **2**. During equilibration, the proteins comprising the capsid segments moved closer to each other in all the systems, and the flexible coils that make contacts between neighboring pentamers rearranged from their initial conformations. The change of the two-pentamer system from the initial to a more compact state is shown in **Figure 1b**. In systems where NPs were present, pentamers also moved closer to each other, while the NPs diffused over the initial 7-9 ns from the aqueous solution towards surfaces of the capsid segments, where they remained bound for the rest of the simulations (**Supplementary Movie**).

The primary binding mode between NP and capsid segments, as observed in the simulations, is shown in **Figures 1c** and **2b,d**. In all the cases, the 2.4 nm gold core NP interacts with the capsid segment at the interface of two L1 pentamers, and over time, it is observed to wedge in between two pentamers that it binds. The NP binds to the interface of two pentamers even in three- and four-pentamer capsid segments, in which there are interfaces of three pentamers within the segments (**Figure 2**). Over the course of the three- and four-pentamer systems simulations, the NPs shifts from three pentamer junctions to two pentamer junctions, and eventually bind to A-F pentamer junction in three-pentamer system (after ∼30 ns) and B-F pentamer junction in four-pentamer system (after ∼70 ns), as shown in **Figure 2b,d**. The binding mode of NP to the capsid segment is likely determined by the NP size: larger NPs may have different binding modes, potentially including the junctions of three pentamers.

**Figure 2.**
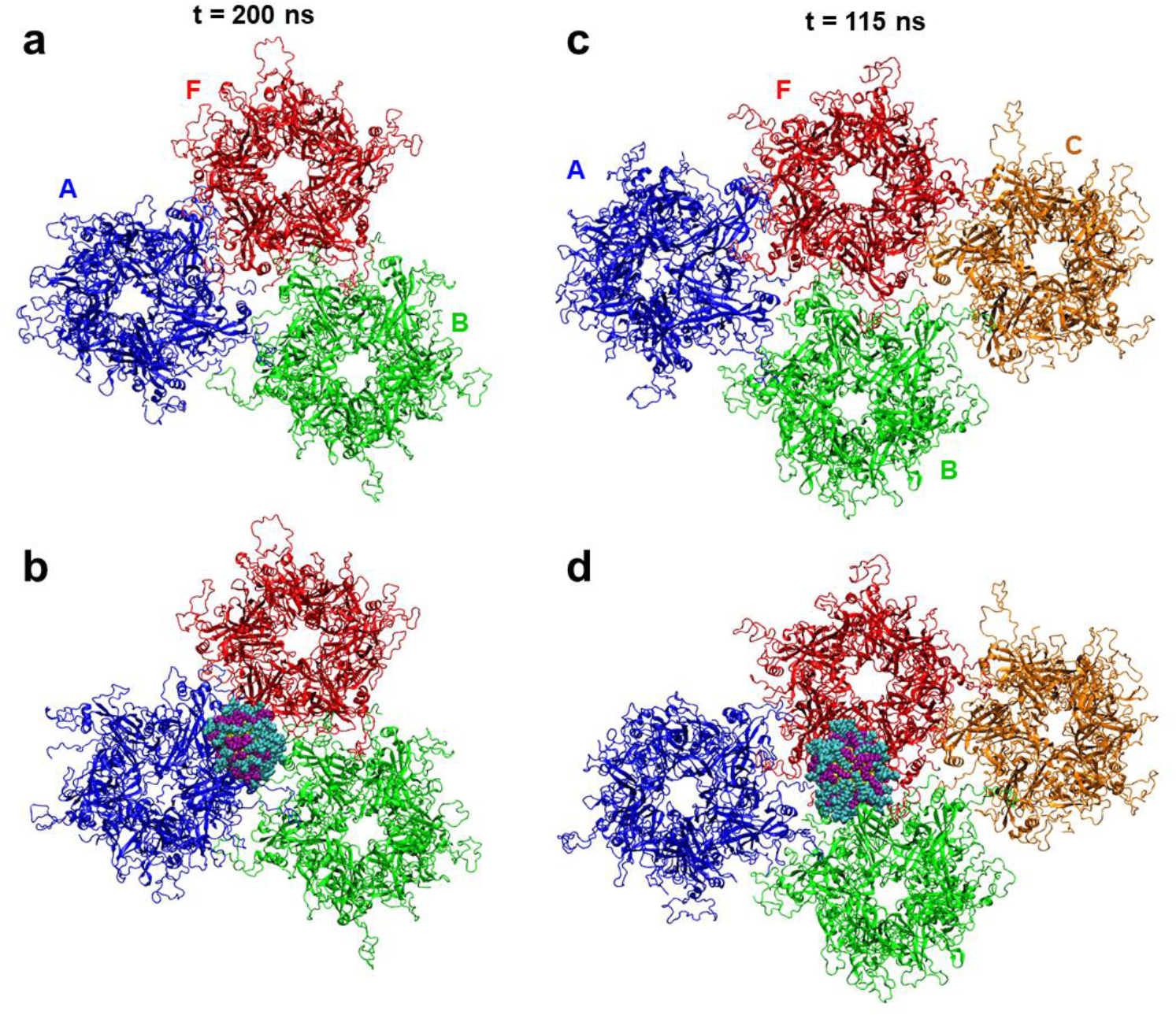
Simulated systems of larger HPV16 capsid segments alone and interacting with sulfonated ligand-coated gold core nanoparticle. a) Conformations of the capsid segment containing three L1 pentamers after 200 ns of MD simulations. b) Conformations of the three pentamer segment in the presence of the MUS:OT NP after 200 ns of MD simulations. c) Conformations of the capsid segment containing four L1 pentamers after 115 ns of MD simulations. b) Conformations of the four pentamer segment in the presence of the MUS:OT NP after 115 ns of MD simulations. The pentamers are colored in blue (A), red (F), green (B), and orange (C), and the aqueous solution is not shown for clarity. The color scheme of the NP is the same as in Figure 1.

In all the examined systems, internal stabilities of individual L1 proteins were tracked by calculating root mean square displacements (RMSDs) of parts with defined secondary structures from the systems simulated without and with the NPs. As **Figures S2** and **S3** show, there are no significant differences in RMSDs of analogous L1 proteins in the systems simulated without and with the NPs. Therefore, the internal stabilities of L1 proteins are not compromised in the presence of NPs.

### Effects of Nanoparticle Binding on the Positions of Proteins within Capsid Segments

MD trajectories of all the systems, both without and with NPs, clearly show that L1 proteins move with respect to each other. To measure the changes in the positions of proteins within capsid segments, we tracked distances between centers of mass (COM) of the adjacent pentamer pairs and angles between the central axes of the adjacent pentamer pairs. The resulting distances and angles between adjacent proteins in two- and three-pentamer systems are shown in **Figure 3**. In two-pentamer systems, distances between A and F pentamers decrease over time from their initial values, both for the lone capsid segment and the capsid segment binding to the NP (**Figure3b**), in agreement with the visual observations that capsid proteins move closer to each other over time. However, these distances decrease by different amounts in the absence and presence of NPs. Namely, the pentamers equilibrate at COM distances of ∼99 Å and ∼ 104 Å in the absence and presence of NP, respectively. Therefore, NP binding to two-pentamer segment leads to an increase in COM distance of ∼ 5 Å between the adjacent pentamers. The pentamers in two-pentamer systems also equilibrate at angles of ∼20° and ∼26° in the absence and presence of NP, respectively (**Figure 3c**). Overall, these pentamers maintain an angle that is ∼ 6° larger on average in the presence of the NP than in its absence. The observations indicate that NP acts as a wedge between the pentamers.

**Figure 3.**
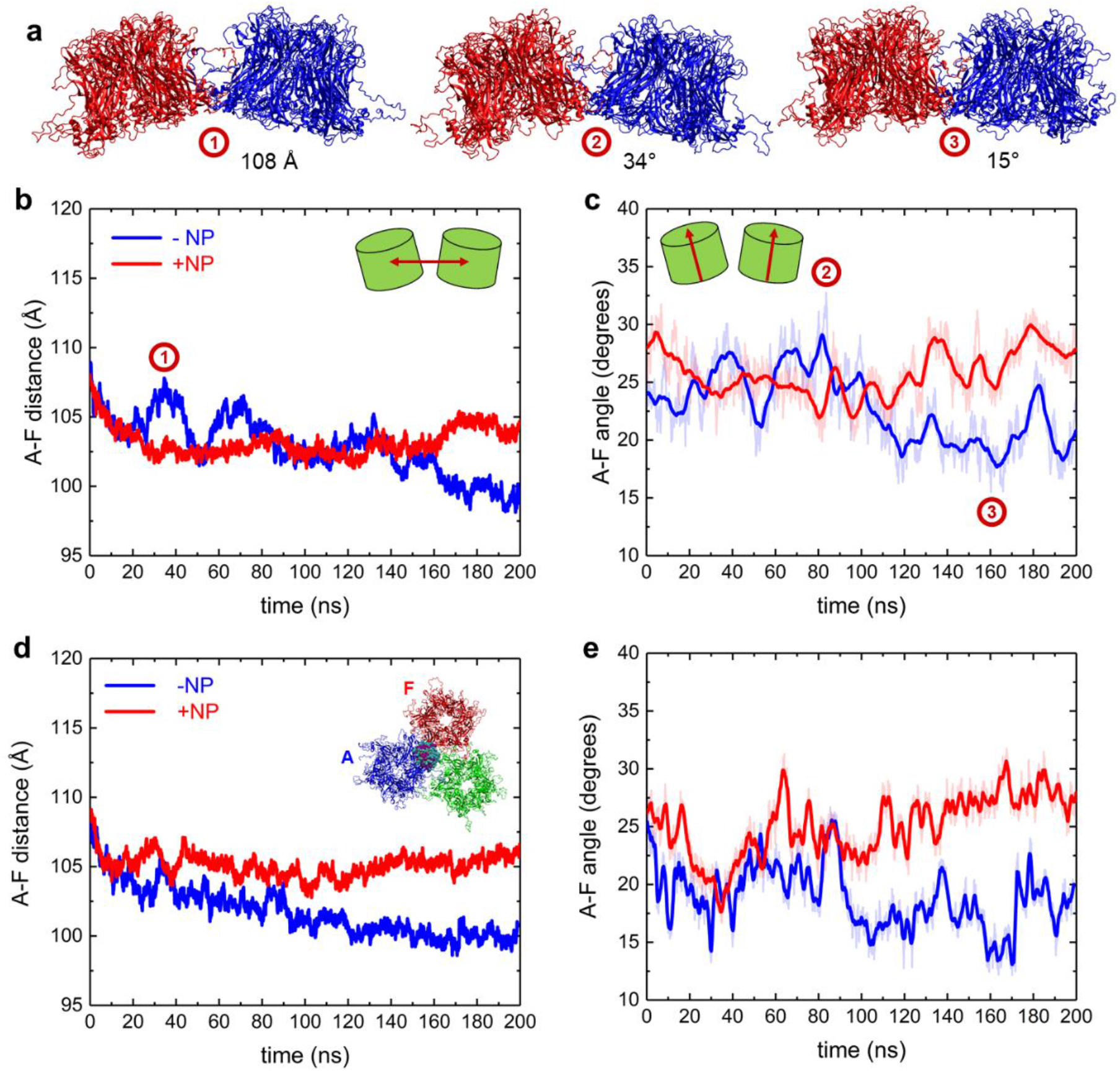
Distances and angles between pentamers in HPV capsid segments alone and when interacting with MUS:OT NP. a) Representative snapshots of two-pentamer capsid segments with pentamers assuming different distances and angles with respect to each other. The snapshot times are labeled in the distance and angle plots in panels (b-c) below. b) Distance between A and F pentamers alone and when interacting with NP for two-pentamer segment systems. c) Angle between A and F pentamers alone and when interacting with NP for two-pentamer segment systems. d) Distance between A and F pentamers alone and when interacting with NP for three-pentamer segment systems. e) Angle between A and F pentamers alone and when interacting with NP for three-pentamer segment systems.

The effects of the NP on protein positions are similar in two- and three-pentamer systems. The distances and the angles between A and F pentamers are on average greater by ∼ 6 Å and ∼ 10°, respectively, in the system with the NP than in the system without the NP. In three-pentamer systems, distances between pentamers whose interfaces do not participate in NP binding are similar in systems with and without the NP (**Figure S4**). For the four-pentamer system, the NP has no clear effects on the segment perturbation, as shown in **Figures S5-S6**. These negligible effects are likely due to a combination of short simulation timescales and a larger network of interactions between the pentamers within the extended capsid that may increase the times over which the segment changes occur.

### Effects of Nanoparticle Binding on the Pentamer Interface and the Integrity of Capsid Segments

When the NP binds at the interface of two pentamers, it interacts with two out of five chains on each pentamer. These chains on A and F pentamers, labeled as A1, A5, F1 and F2, are highlighted in **Figure 4a**. Similar interactions are observed for NPs within three- and four-pentamer systems. The individual chains preserve their secondary structures while interacting with NPs (**Figures S7-S9**), as it is mostly the flexible loop regions of the chains, protruding from the capsid surface into the solvent, that interact with NPs (**Figures 4b, S10**). These loop regions and specifically some of the lysine residues on them (K278, K356, K361, K54 and K59) were found to be implicated in the recognition of heparan sulfate proteoglycans^24,34^, indicating that NPs coated with MUS and OT ligands can mimic HSPG cell receptors allowing for effective viral association.

**Figure 4.**
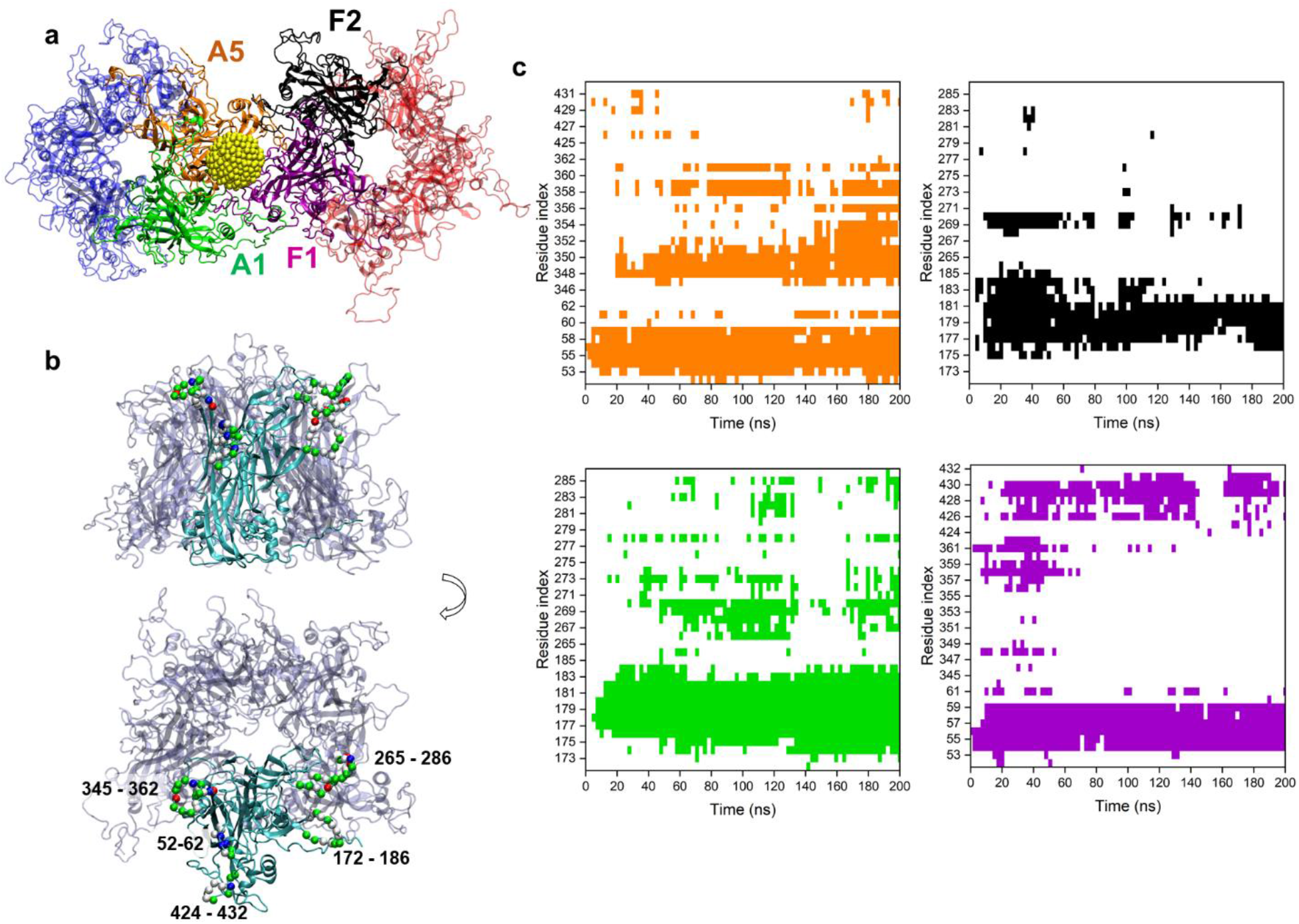
Interactions of MUS:OT NP at the interface of two L1 pentamers in the two-pentamer system. a) A snapshot of the NP interacting at the interface of two pentamers, shown in faded blue and red. The NP interacts with two distinct chains of both A and F pentamers, labeled as A1 (green), A5 (orange), F1 (purple) and F2 (black). The NP gold core is shown in yellow, and MUS and OT ligands are not shown for clarity. b) Side and top views of two chains of the pentamer typically found to interact with the NP, highlighted in red. The residues consistently found to interact with NP are shown by their Cα atoms (green, white, blue and red spheres label their residue types, namely polar, non-polar, basic and acidic) and labeled with residue indices. c) Contact maps of residues of the four chains of two-pentamer systems found to interact with NP over the course of the simulation. Colors in the contact maps match the colors of the four chains labeled within pentamers in panel a (A5 – orange, F2 – black, A1 – green, and F1 – purple). White regions in contact maps denote no contact.

As each chain is equivalent in sequence, similar in structure, but different in orientation, MUS:OT NP interacts with two unique surfaces of A1, A5, F1 and F2 chains. **Figures 4b** and **S10** highlight the residues of these chains that interact significantly with the NP: chains F1 and A5 interact with NP via three loop regions (residues 52-62, 345-362 and 424-432), whereas chains A1 and F2 interact with NP via two loop regions (residues 172-186 and 265-286). Contacts between these loop regions and the NP are largely maintained over the simulation time, as shown in the contact maps in **Figure 4c**. These loop regions contain many charged and non-polar amino acids (**Figure S10**). For example, the loop region with residues 52-62 interacts strongly with the NP via K53, K54, and K59, as well as the polar N56-58 residues. Interactions of NP with this loop region (residues 52-62) is significant in all three examined systems indicating the importance of this region for establishing the initial NP-capsid binding through long range electrostatic interactions and preserving it later via local electrostatic and H-bond interactions. All the loops, and especially the loop with residues 172-186, also interact with the NP via its hydrophobic residues. The interactions of NP with capsid segments via charge-charge and hydrophobic interactions correlates with observations of a previous MD study of a single L1 pentamer with MUS:OT NP. The charge-charge interactions occur between negative sulfonate groups on MUS:OT-NP and the positive HSPG-binding lysine residues of loop regions, while the hydrophobic interactions occur between non-polar alkyl chains of NP ligands and nonpolar groups on L1 protein surface. Similar pentamer-NP contacts are observed in the three-pentamer system as in the two-pentamer system (**Figure S11**). However, significantly fewer pentamer-NP contacts are observed in the four-pentamer system (**Figure S12**), indicating a need for longer simulation times in this system to capture the effects of the NP presence on the capsid segment integrity.

We next examine in **Figure 5** how the contacts between neighboring pentamers evolve over time in the absence and presence of the NP within two-pentamer systems. Most of the time, the same inter-pentamer contacts are observed in the absence and presence of the NP. However, towards the end of the 240 ns trajectories, some contacts between pentamers become less frequent or are lost in the presence of the NP, such as the contacts that A5 segment makes with the F pentamer (residues 52 – 62 and amino acids surrounding the residue 449) and the contacts that F2 segment makes with the A pentamer (residues 169 – 189). Similar loss of interactions between pentamers interacting with the NP is observed in three-pentamer and four-pentamer systems (although less pronounced, as shown **Figures S13** and **S14**). All these contact maps indicate that NPs are able to disrupt some interactions between neighboring pentamers over the simulation timescales.

**Figure 5.**
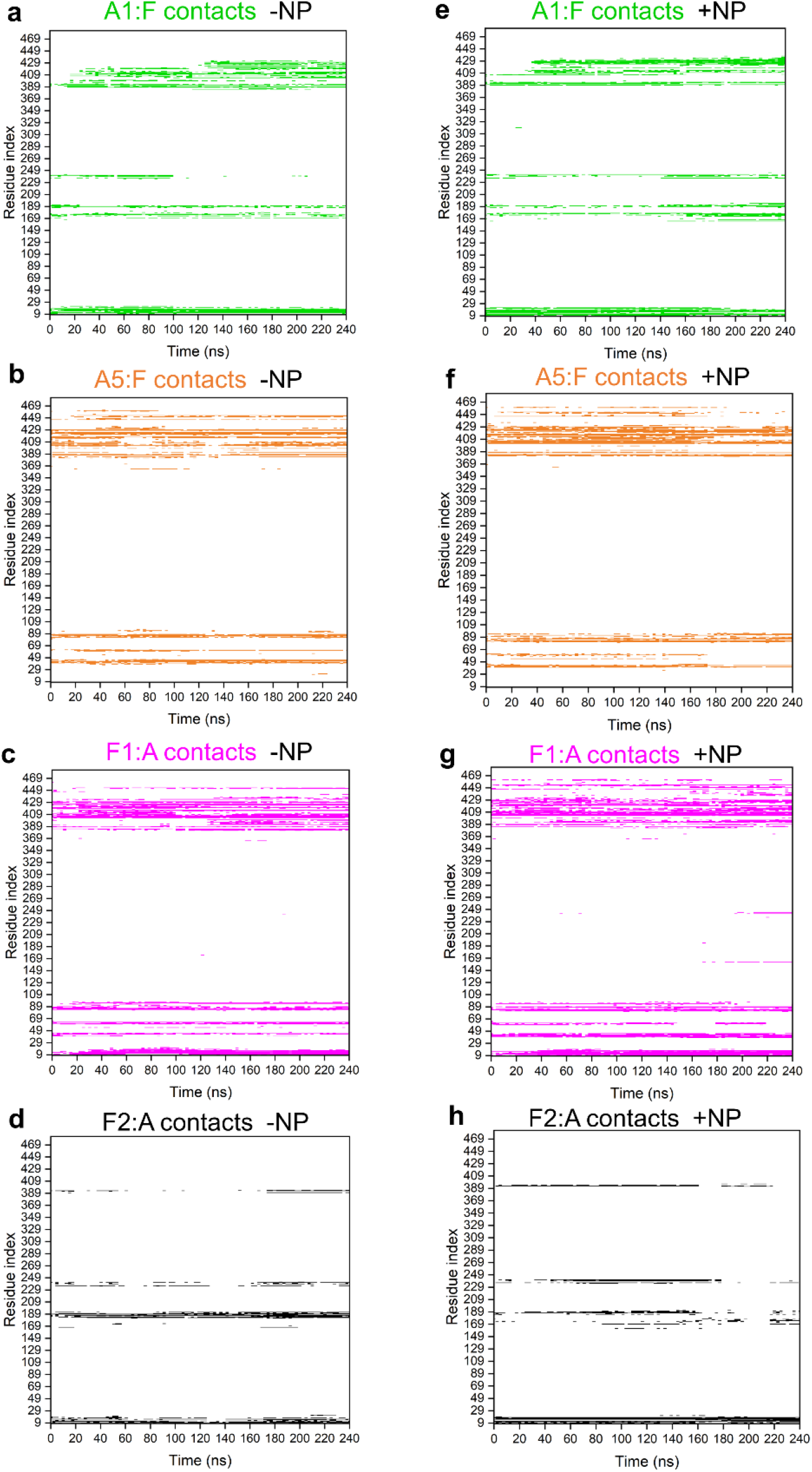
Contact maps of L1 pentamer interactions in two-pentamer segments alone and when interacting with MUS:OT NP. a-d) Contact maps of A1 and A5 chain residues in contact with F pentamer and F1 and F2 chain residues in contact with A pentamer within two-pentamer systems over the course of the simulation. e-h) Contact maps of A1 and A5 chain residues in contact with F pentamer and F1 and F2 chain residues in contact with A pentamer within two-pentamer systems in the presence of MUS:OT NP over the course of the simulation. The colors in the contact maps match the colors of the four chains within pentamers shown in Figure 4a. White regions in the contact maps denote no contact.

Overall, our studies also show that most of the residues involved in pentamer-NP and pentamer-pentamer interactions are mutually exclusive over the course of 115 – 240 ns simulations. With longer simulation times, more significant effects of NP presence are expected on pentamer-pentamer interactions.

## 4. Conclusion

In this work, we described the interactions between virucidal MUS:OT nanoparticles with 2.4 nm gold cores and segments of HPV16 capsids using atomistic MD simulations. It was previously shown that HSPG-mimicking ligands of these NPs form multivalent interactions with L1 capsid proteins^19^. Here, we determine the initial mechanisms of the virucidal activity and depict it in the scheme of **Figure 6**. The first step of this activity is the binding of MUS:OT NPs at interfaces of two L1 proteins forming the capsid segments. The insertion of the NP at the interface of two L1 proteins leads to increased distances and increased angles between these neighboring proteins. As the time progresses, the NP presence can lead to breaking of some contacts between two neighboring proteins, although our simulations captured the very initial stages of this stage of the virucidal activity. Therefore, the disruption of the HPV16 capsid by MUS:OT NPs is expected to arise at interfaces of two L1 capsid proteins. The revealed mechanism can be utilized for designing new generations of materials that can perturb and disintegrate viral capsids^35^ or control the integrity of other naturally occurring or engineered protein assemblies^36,37^.

**Figure 6.**
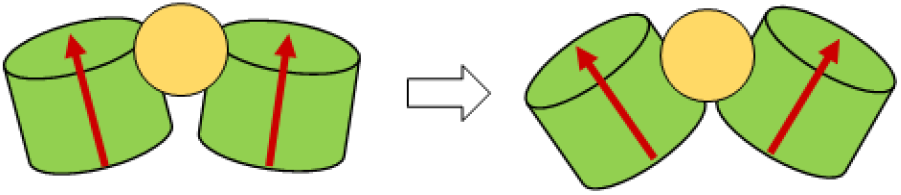
A scheme of MUS:OT NP effect on HPV16 capsid segments. MUS:OT NPs (2.4 nm gold cores) bind to interfaces of two L1 proteins and wedge in between them. The NPs eventually weaken and break the interactions between L1 proteins at this interface.

## Acknowledgment

Research reported in this publication was supported by the National Institute of Allergy and Infectious Diseases of the National Institutes of Health under Award Number R03AI142553. The content is solely the responsibility of the authors and does not necessarily represent the official views of the National Institutes of Health. The authors gratefully acknowledge computer time provided by the Texas Advanced Computing Center (TACC).

## Notes

### Competing Interest Statement

The authors have declared no competing interest.

