## Supplementary Information for "Computational modeling of virucidal inhibition mechanism for broad-spectrum antiviral nanoparticles and HPV16 capsid segments"

Parth Chaturvedi, Payam Kelich, Tara A. Nitka, Lela Vuković*
Department of Chemistry and Biochemistry, University of Texas at El Paso, El Paso, TX 79968, USA

*

Electronic Supporting Information (ESI)

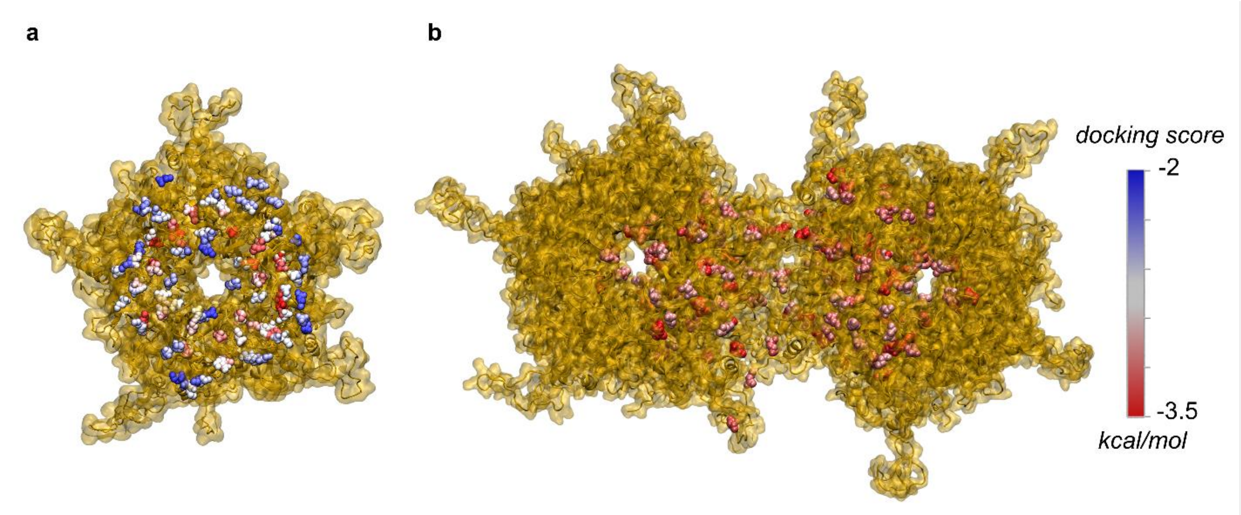


**Figure S1.** The most favorable positions of sulfonated ligands ($CH_3CH_2SO_3^-$) on the HPV16 L1 capsid protein surface determined by docking calculations. a) Structures of ligands docked on the surface of a single L1 protein, colored according to the docking score. b) Structures of ligands docked on the surface of a pair of L1 proteins, colored according to the docking score. In this panel, only the ligands with the lowest docking scores are shown.

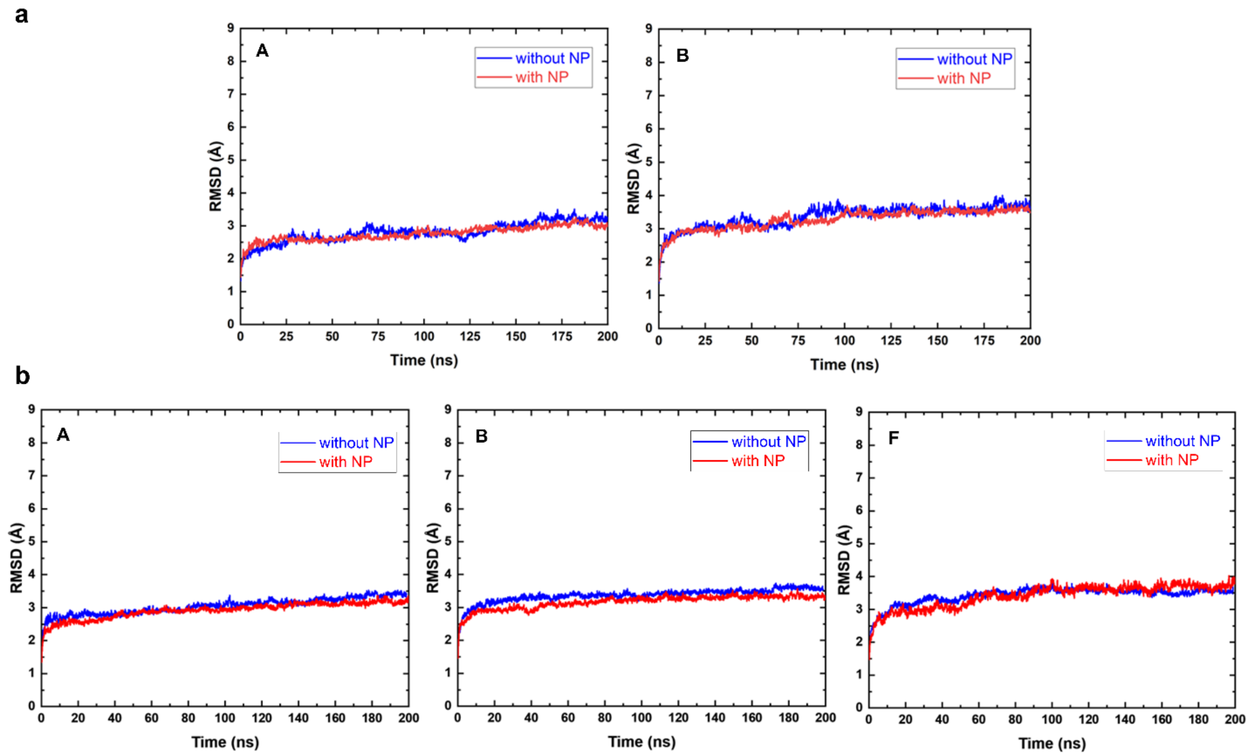


**Figure S2.** RMSDs of L1 pentamers (parts with defined secondary structures, consisting of α-helices and β-sheets) observed during MD simulations for: a) two-pentamer systems, and b) three-pentamer systems.

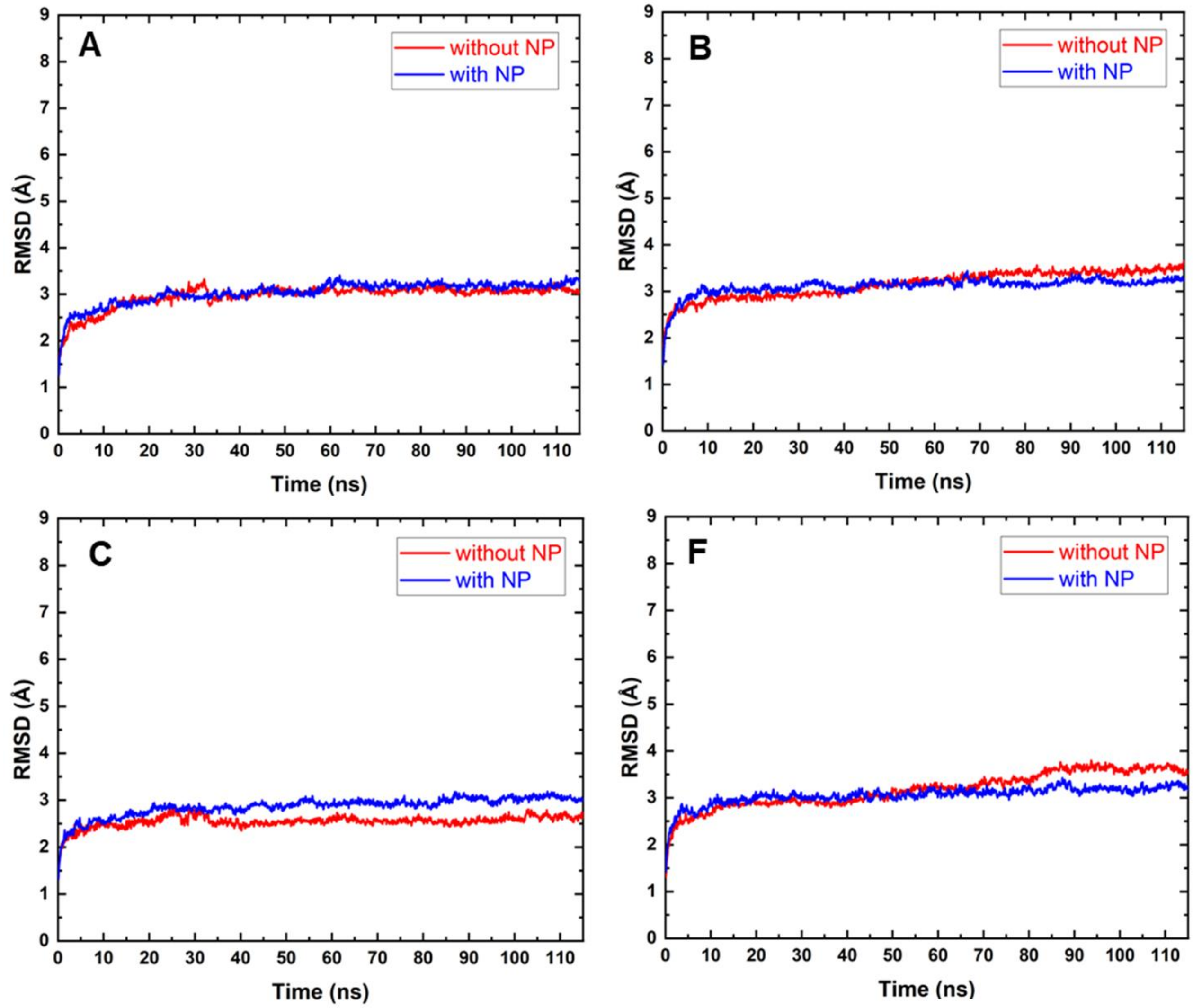


**Figure S3.** RMSDs of L1 pentamers (parts with defined secondary structures, consisting of α-helices and β-sheets) observed during MD simulations for four-pentamer systems.

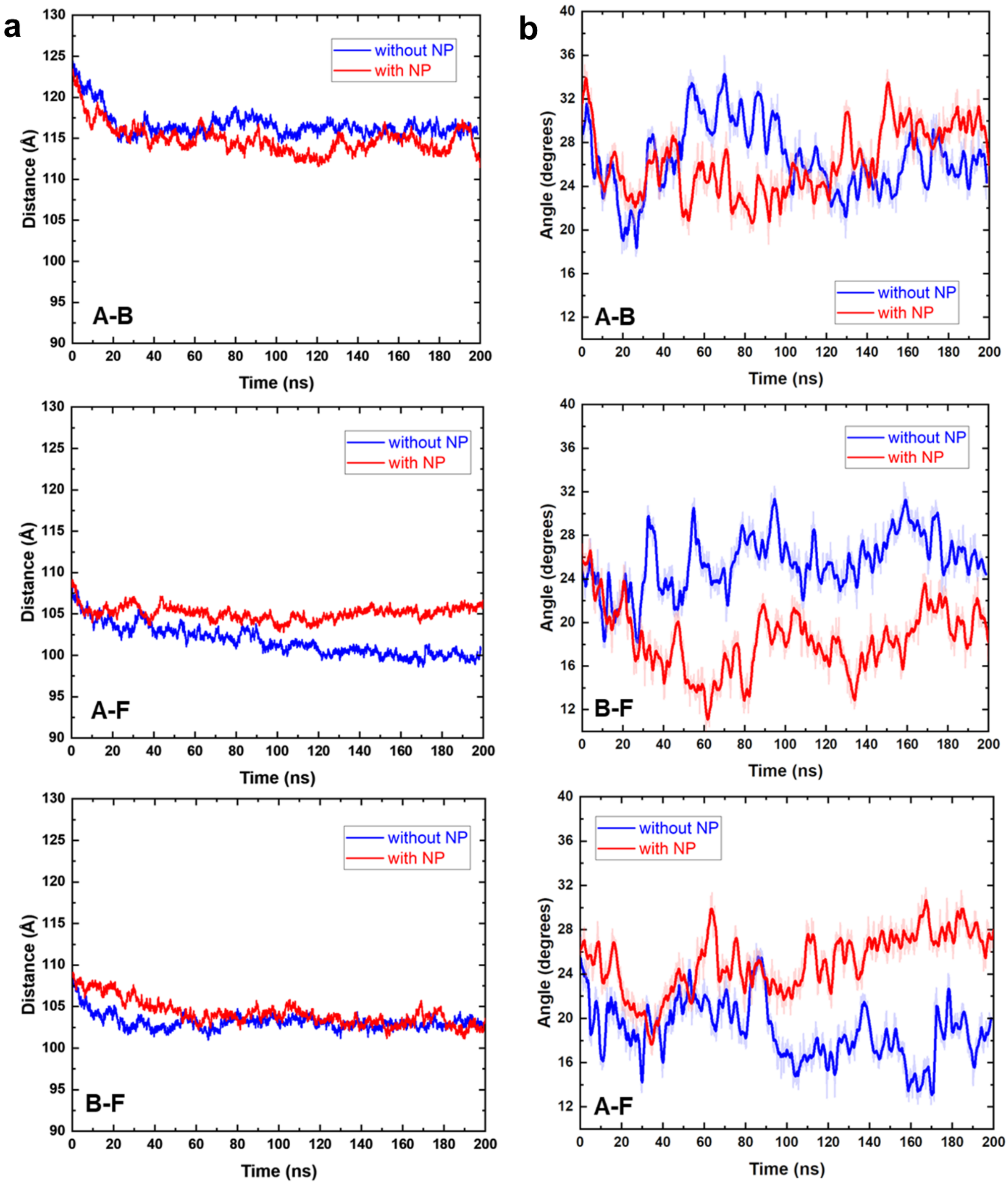


**Figure S4**. a) Distances between centers of mass (COMs) of all two L1 pentamer pairs in three-pentamer systems observed during MD simulations. b) Angles between all two L1 pentamer pairs in three-pentamer systems observed during MD simulations.

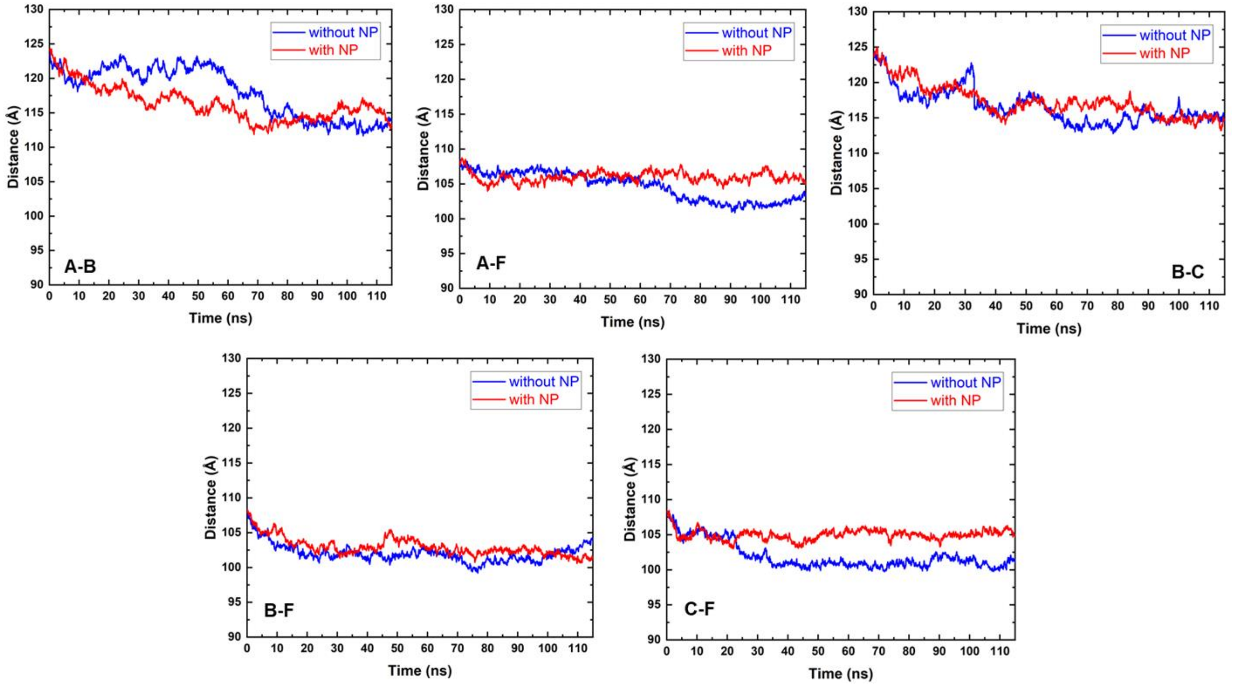


**Figure S5**. Distances between COMs of all two L1 pentamer pairs in four-pentamer systems observed during MD simulations.

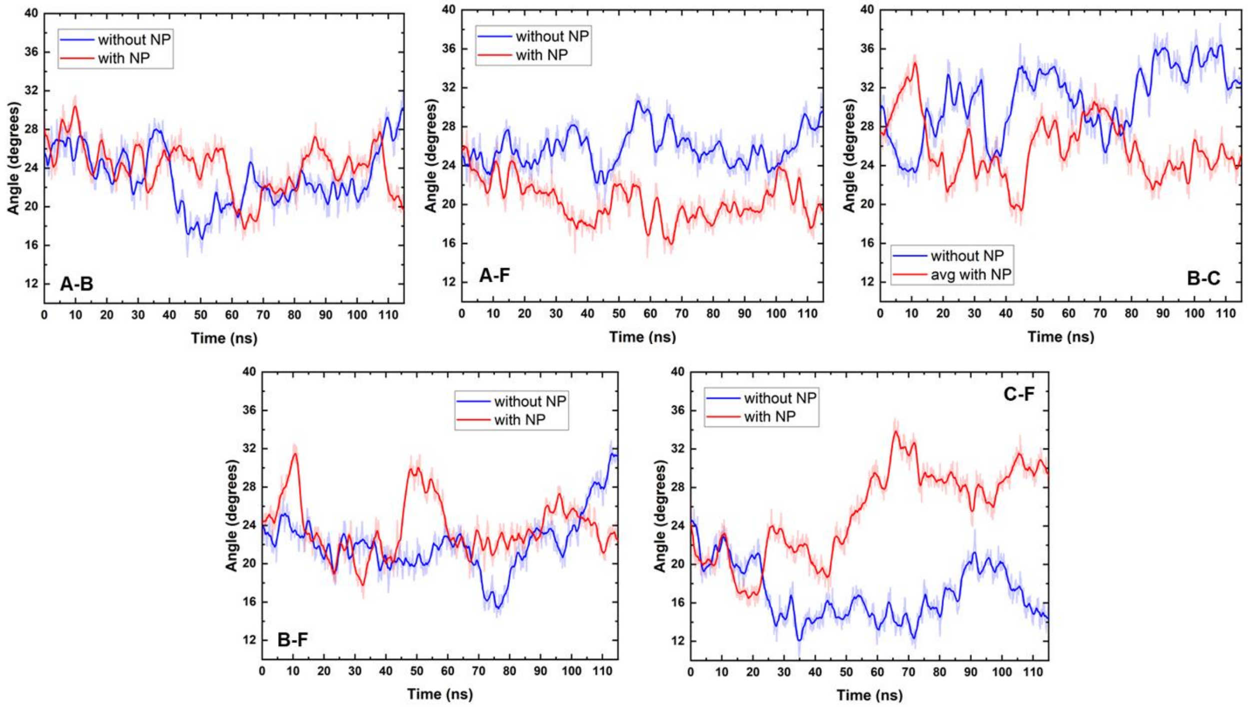


**Figure S6**. Angles between all two L1 pentamer pairs in four-pentamer systems observed during MD simulations.

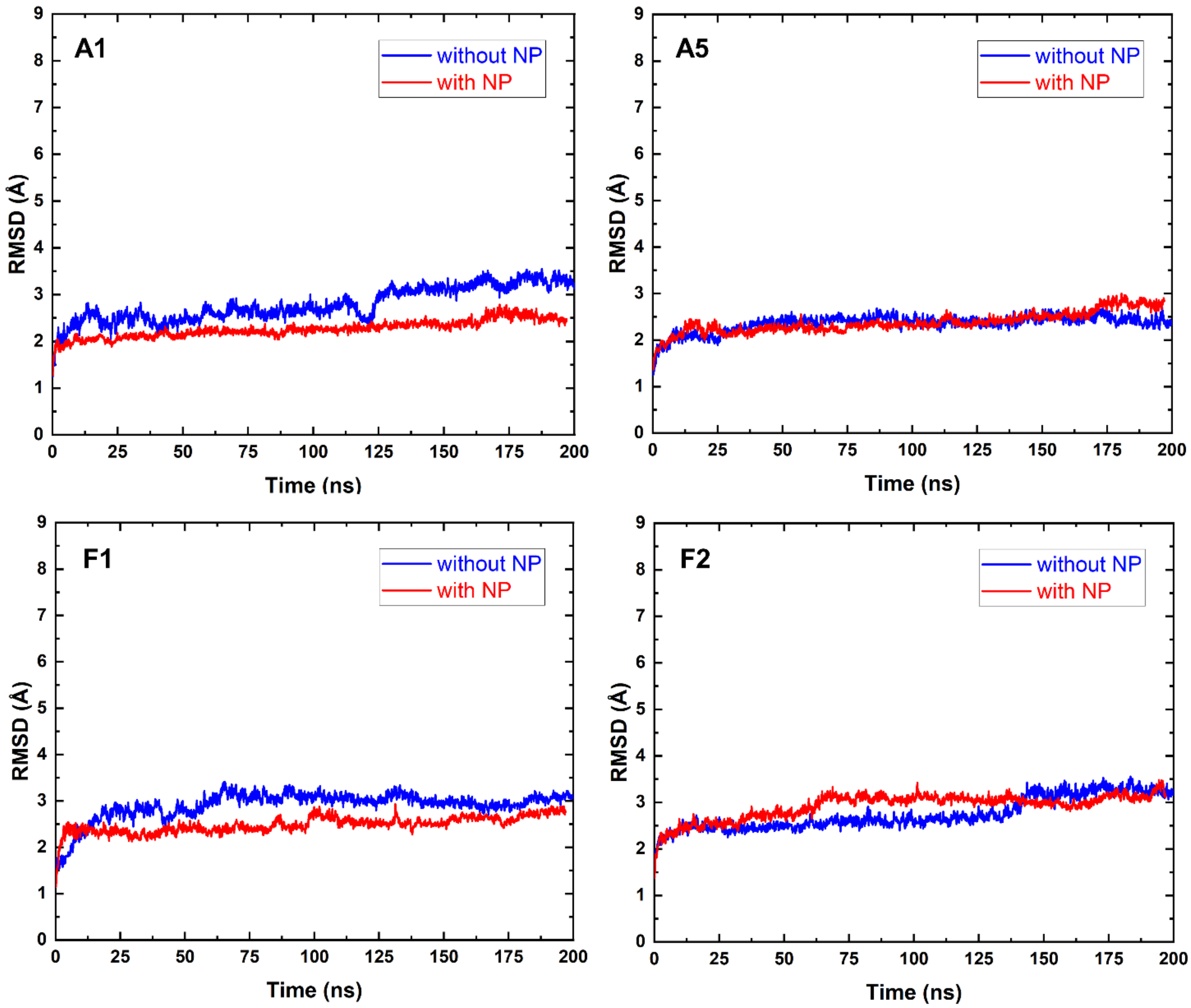


**Figure S7.** RMSDs of four NP-binding chains (their parts with defined secondary structures, consisting of α-helices and β-sheets) within two-pentamer systems observed during MD simulations.

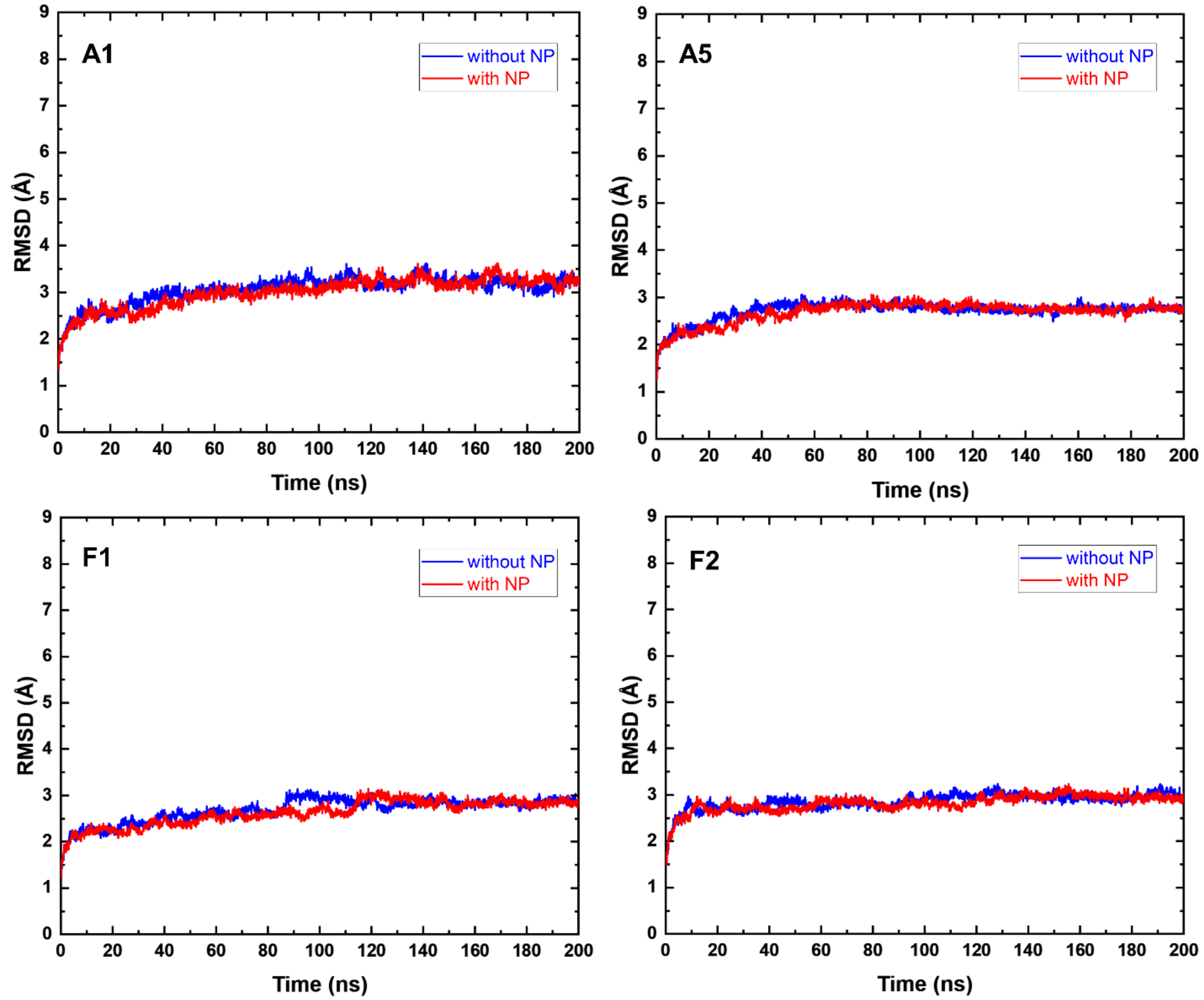


**Figure S8.** RMSDs of four NP-binding chains (their parts with defined secondary structures, consisting of α-helices and β-sheets) within three-pentamer systems observed during MD simulations.

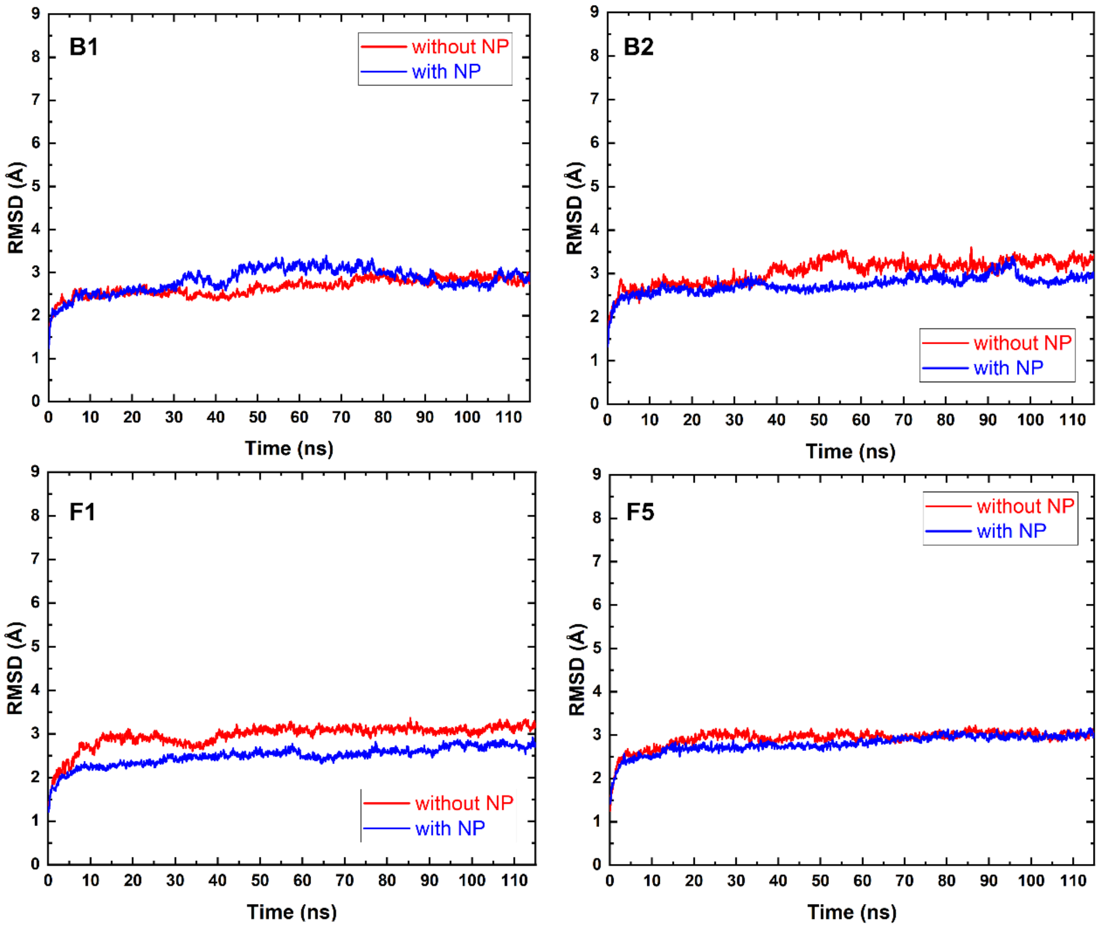


**Figure S9.** RMSDs of four NP-binding chains (their parts with defined secondary structures, consisting of α-helices and β-sheets) within four-pentamer systems observed during MD simulations.

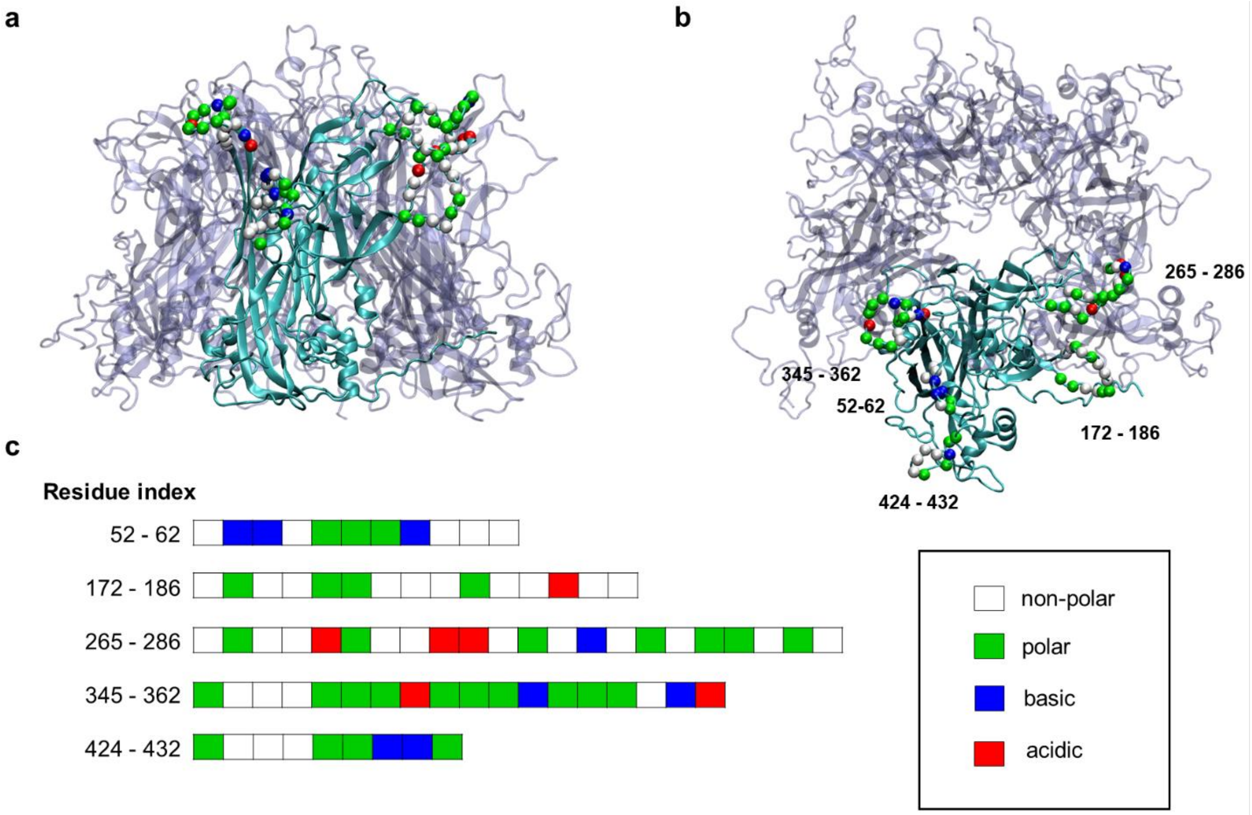


**Figure S10.** a) Side view and b) top view of two typical segments (cyan) of the pentamer with residues interacting with NP, shown as spheres that are colored according to their residue types. c) Residues interacting with the NP shown according to their respective types.

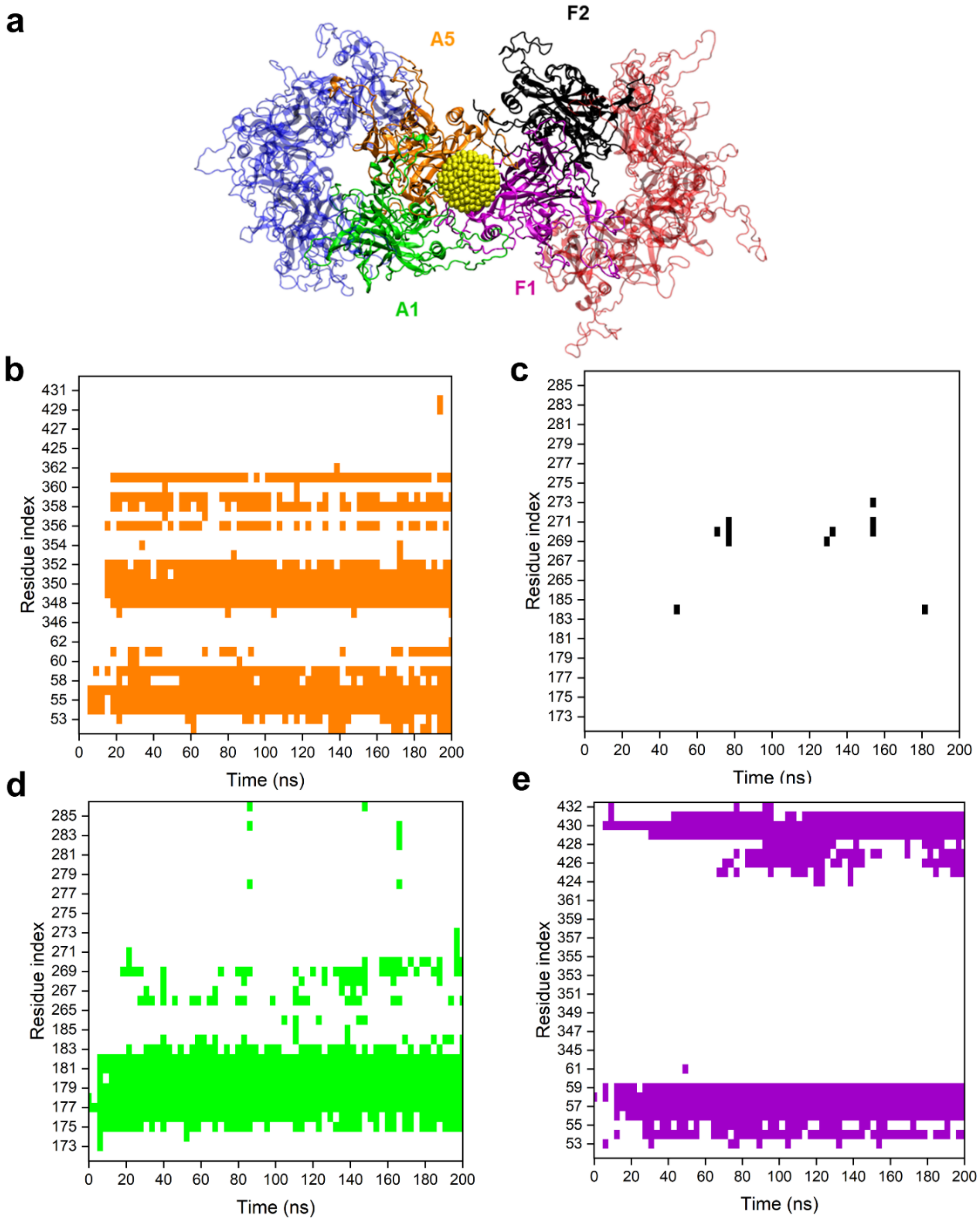


**Figure S11.** Interactions of MUS:OT NP at the interface of two L1 pentamers in the three-pentamer system. a) A snapshot of the NP interacting at the interface of two pentamers, shown in faded blue and red. The NP interacts with two distinct chains of both A and F pentamers, labeled as A1 (green), A5 (orange), F1 (purple) and F2 (black). The NP gold core is shown in yellow, and MUS and OT ligands are not shown for clarity. b-e) Contact maps of residues of the four chains of three-pentamer systems found to interact with NP over the course of the simulation. Colors in the contact maps match the colors of the four chains labeled within pentamers in panel a (A5 – orange, F2 – black, A1 – green, and F1 – purple). White regions in contact maps denote no contact.

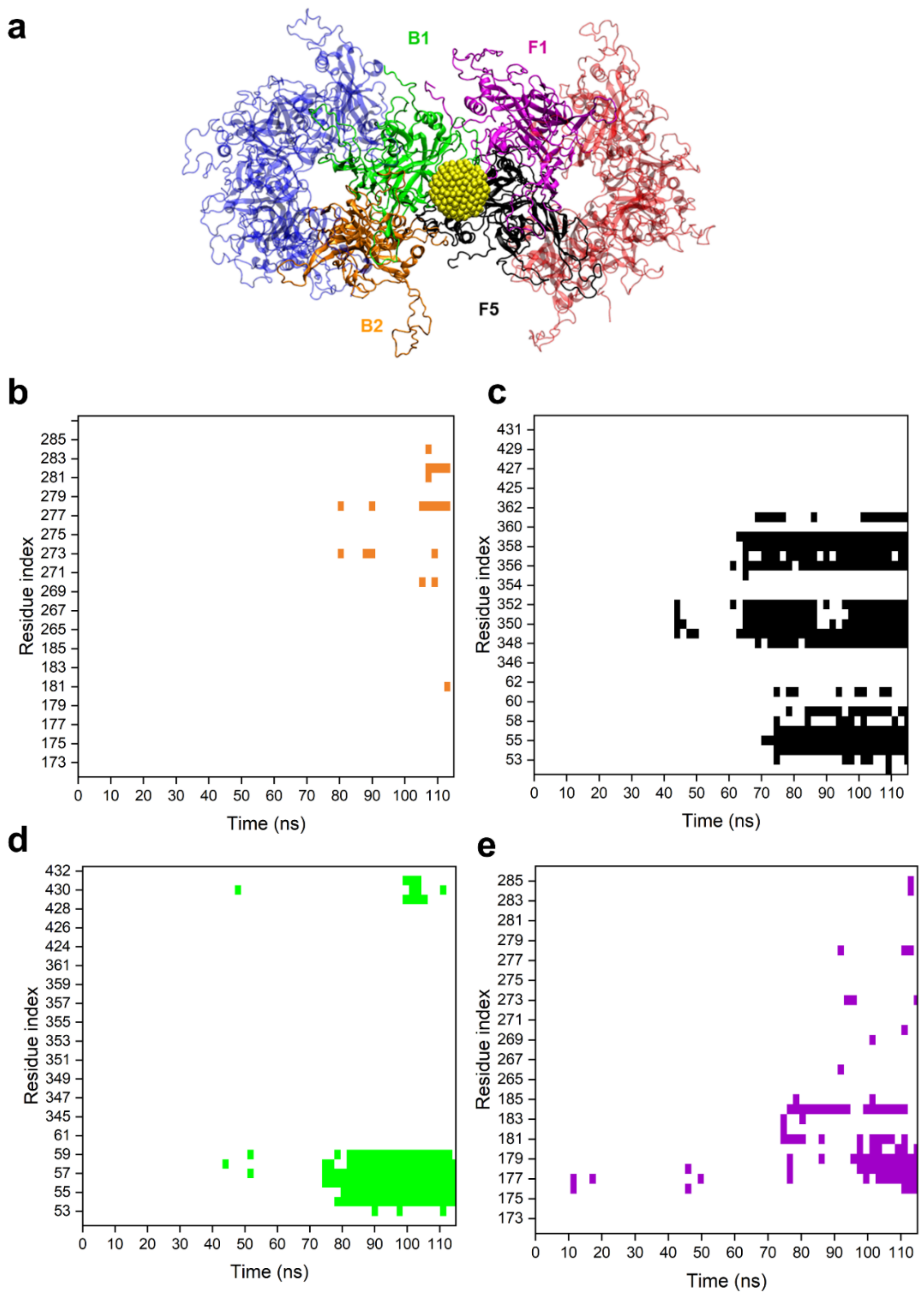


**Figure S12.** Interactions of MUS:OT NP at the interface of two L1 pentamers in the four-pentamer system. a) A snapshot of the NP interacting at the interface of two pentamers, shown in faded blue and red. The NP interacts with two distinct chains of both B and F pentamers, labeled as B1 (green), B2 (orange), F1 (purple) and F5 (black). The NP gold core is shown in yellow, and MUS and OT ligands are not shown for clarity. b-e) Contact maps of residues of the four chains of four-pentamer systems found to interact with NP over the course of the simulation. Colors in the contact maps match the colors of the four chains labeled within pentamers in panel a. White regions in contact maps denote no contact.

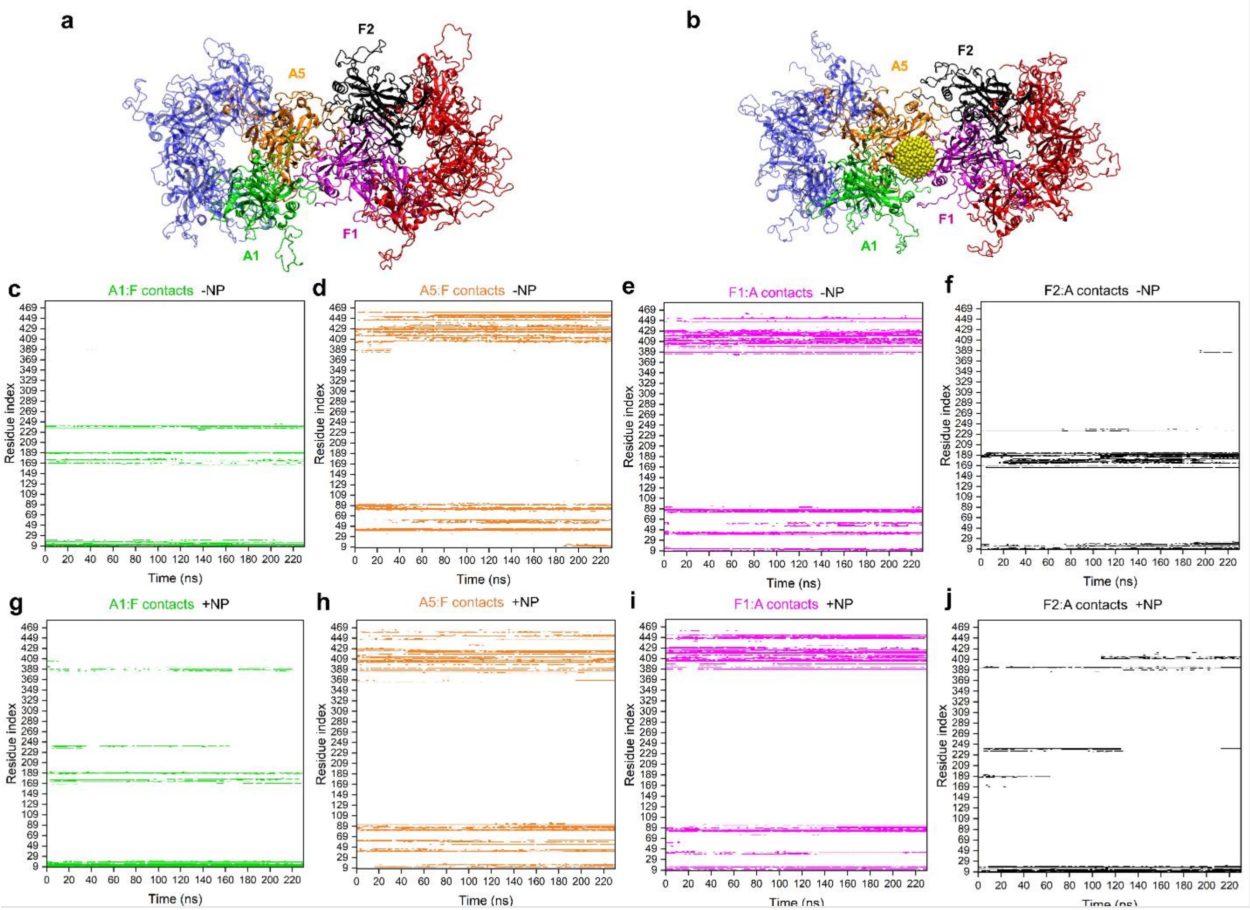


**Figure S13.** a) A snapshot of the two interacting pentamers in three-pentamer system. b) A snapshot of the NP interacting at the interface of the same two pentamers within the three-pentamer system. Chains of neighboring pentamers interacting with each other are labeled in distinct colors. The NP gold core is shown in yellow, and MUS and OT ligands are not shown for clarity. c-f) ) Contact maps of A1 and A5 chain residues in contact with F pentamer and F1 and F2 chain residues in contact with A pentamer within three-pentamer systems in the absence of MUS:OT NP over the course of the simulation. g-j) Contact maps of A1 and A5 chain residues in contact with F pentamer and F1 and F2 chain residues in contact with A pentamer within four-pentamer systems in the presence of MUS:OT NP over the course of the simulation.

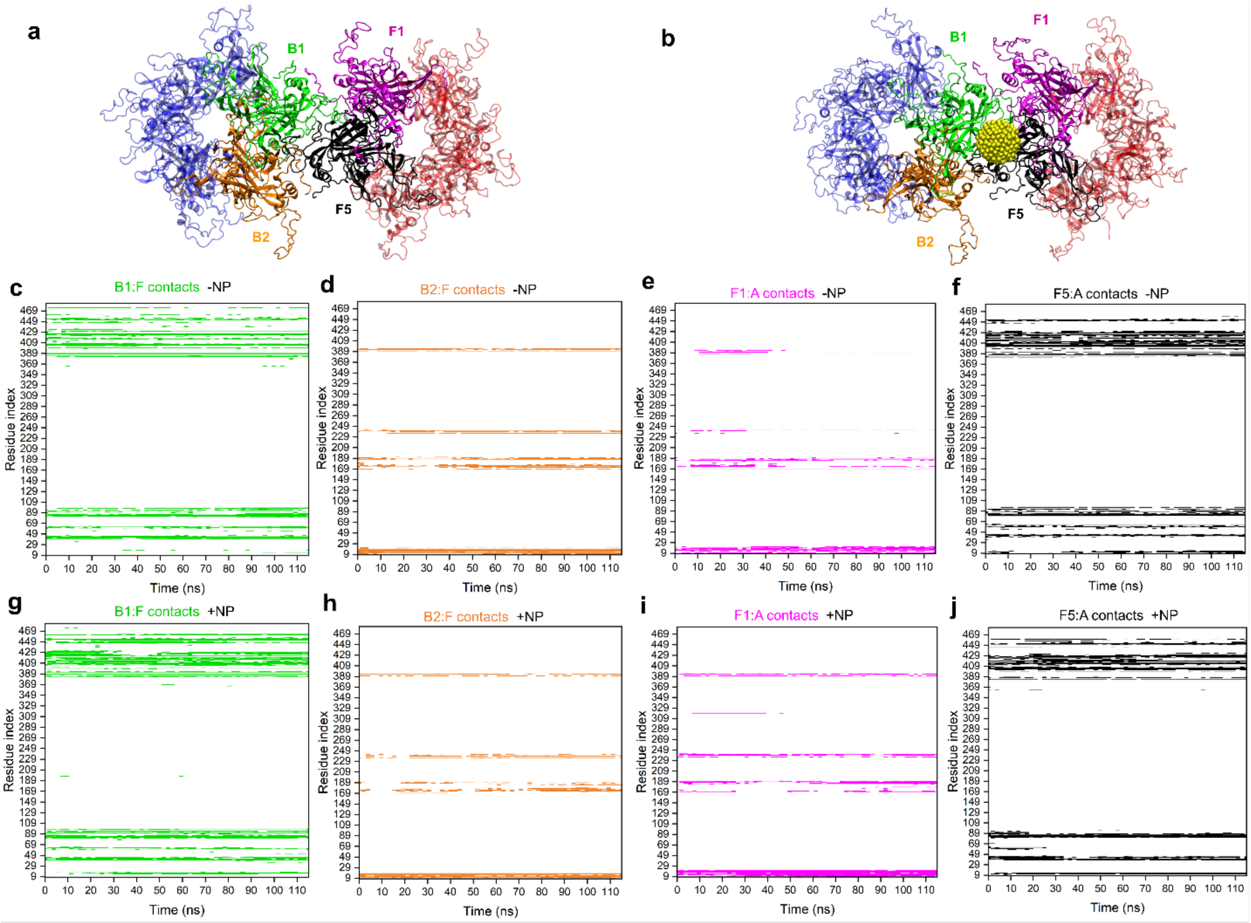


**Figure S14.** a) A snapshot of two interacting pentamers in the four-pentamer system. b) A snapshot of the NP interacting at the interface of the same two pentamers within the four-pentamer system. Chains of neighboring pentamers interacting with each other are labeled in distinct colors. The NP gold core is shown in yellow, and MUS and OT ligands are not shown for clarity. c-f) ) Contact maps of B1 and B2 chain residues in contact with F pentamer and F1 and F5 chain residues in contact with A pentamer within four-pentamer systems in the absence of MUS:OT NP over the course of the simulation. g-j) Contact maps of B1 and B2 chain residues in contact with F pentamer and F1 and F5 chain residues in contact with B pentamer within four-pentamer systems in the presence of MUS:OT NP over the course of the simulation.
